# DeepREAL: A Deep Learning Powered Multi-scale Modeling Framework Towards Predicting Out-of-distribution Receptor Activity of Ligand Binding

**DOI:** 10.1101/2021.09.12.460001

**Authors:** Tian Cai, Kyra Alyssa Abbu, Yang Liu, Lei Xie

**Affiliations:** Ph.D. Program in Computer Science, The Graduate Center, The City University of New York, New York, 10016, USA; Department of Computer Science, Hunter College, The City University of New York, New York, 10065, USA; Helen and Robert Appel Alzheimer’s Disease Research Institute, Feil Family Brain & Mind Research Institute, Weill Cornell Medicine, Cornell University, New York, 10021, USA

## Abstract

Drug discovery has witnessed intensive exploration of the problem of drug-target physical interactions over two decades, however, a strong drug binding affinity to a single target often fails to translate into desired clinical outcomes. A critical knowledge gap needs to be filled for correlating drug-target interactions with phenotypic responses: predicting the receptor activities or function selectivity upon the ligand binding (i.e., agonist vs. antagonist) on a genome-scale and for novel chemicals. Two major obstacles compound the difficulty on this direction: known data of receptor activity is far too scarce to train a robust model in light of genome-scale applications, and real-world applications need to deploy a model on data from various shifted distributions. To address these challenges, we have developed an end-to-end deep learning framework, DeepREAL, for multi-scale modeling of genome-wide receptor activities of ligand binding. DeepREAL utilizes self-supervised learning on tens of millions of protein sequences and pre-trained binary interaction classification to solve the data distribution shift and data scarcity problems. Extensive benchmark studies that simulate real-world scenarios demonstrate that DeepREAL achieves state-of-the-art performance in out-of-distribution settings.

## 1 Introduction

Over the last two decades, drug discovery has been dominated by target-based high-throughput compound screening. Unfortunately, this “one-drug-one-gene” approach has been costly and had a low success rate due to our limited understanding of molecular and cellular mechanisms of drug actions[4][22]. Drugs from the target-based screening often interact with unexpected off-targets, leading to serious side effects[13][16]. Furthermore, a polypharmacology approach is often needed to achieve desired therapeutic efficacy and overcome drug resistance for multi-genic diseases[23]. In order to predict drug phenotypic response at the organismal level, it is necessary to not only elucidate genome-scale drug-target interactions (DTIs) but also reveal how DTIs collectively modulate a biological system.

The drug mode of action is a multi-scale process that starts with drug binding to its targets, principally proteins. Then the drug can act as an antagonist or an agonist to block or enhance downstream biological processes, respectively. Therefore, it is critically important to model the change of receptor activities or functional selectivity upon the drug binding for understanding how the drug modulates physiological functions. The information on the receptor activity following the ligand binding will fill in a critical knowledge gap in correlating DTIs to clinical outcomes. Although a great deal of efforts have been devoted to predict genome-wide DTIs using deep learning[2], few large-scale experimental and computational studies have been able to specify the receptor activity of ligand binding, i.e., the functional selectivity of the ligand as an antagonist or an agonist.

In this research, we aim to predict not only whether any pairs of proteins and chemicals interact with each other or not but also the receptor activity upon the binding, especially, making reliable predictions for understudied “dark” proteins that do not have any ligand annotations[18] and novel chemicals whose structures are different from those in the training data. Recent work [19] applied Random Forest to train an independent machine learning model for each individual Opioid receptor to predict their receptor activity. Unfortunately, labeled data for the receptor activity are scarce. Only a limited number of receptors have sufficient function selectivity data to train a robust and data-hungry deep learning model. Thus, the one-protein-one-model approach cannot be extended to majority of proteins that have few or no labeled data[19]. It is a challenging task to predict the function for dark proteins in general using machine learning. Conventional machine learning methods assume that the distribution of unseen data and training data is identically and independently distributed (IID). This assumption may not hold for the dark proteins that are dissimilar from those in the training data. In other words, many dark proteins are out-of-distribution (OOD) in terms of the training samples. Similarly, unseen novel chemicals are also OOD cases. To address the data scarcity and OOD challenges, we have developed an Artificial Intelligence (AI)-powered multi-scale modeling framework, DeepREAL, to simulate the multi-scale drug actions and predict the receptor activity of ligand bindings for dark proteins and novel chemicals. We first apply self-supervised learning to train a protein sequence model for a universal protein sequence embedding on a genome scale. This allows us to detect subtle relationships between dark proteins and ligand-annotated proteins. We then train a binary classification deep learning model to predict whether a chemical binds to a protein and extract a latent presentation of DTIs. Because there is a large amount of binary interaction data, it is possible to train a robust deep learning model. Finally, we integrate chemical fingerprint model, sequence embedding model, and DTI latent representation model to train an end-to-end deep learning model for predicting the receptor activity of ligand binding using limited data. In the rigorous benchmark studies that simulate real-world applications, DeepREAL significantly improves the generalization ability in the OOD setting compared with the state-of-the-art methods[19][2]. Data and code is available on github https://github.com/XieResearchGroup/DeepREAL.

## 2 Results and Discussion

### 2.1 Overview of methods

Given a chemical structure and the sequence of a receptor protein, DeepREAL will predict whether the chemical is an agonist or an antagonist if it binds to the receptor, or not bind to it at all (Figure 1A). Intuitively, DeepREAL leverages large data sets to hierarchically inform predictions on the receptor activity whose labeled data are scarce along a three-stage end-to-end pipeline as illustrated in Figure 1B. By pretraining protein sequences on whole Pfam in Stage 1, DeepREAL equips itself with genome-scale protein representations that capture novel relationships between proteins beyond sequence homology[2]. By pretraining on a large scale of binary DTI data in Stage 2, DeepREAL builds knowledge of chemical-protein interactions which is the initial step in the ligand binding event. Finally, in Stage 3, information learned from sequence embeddings and binary interactions are transferred into predicting receptor activities using a small amount of data. This hierarchy design maintains knowledge learned from heterogeneous resources and enhance model robustness in the face of shifted data distribution during deployment. More details of DeepREAL design and implementation could be found in Methods section. Detailed model architecture is in Supporting Figure S1.

**Figure 1:**
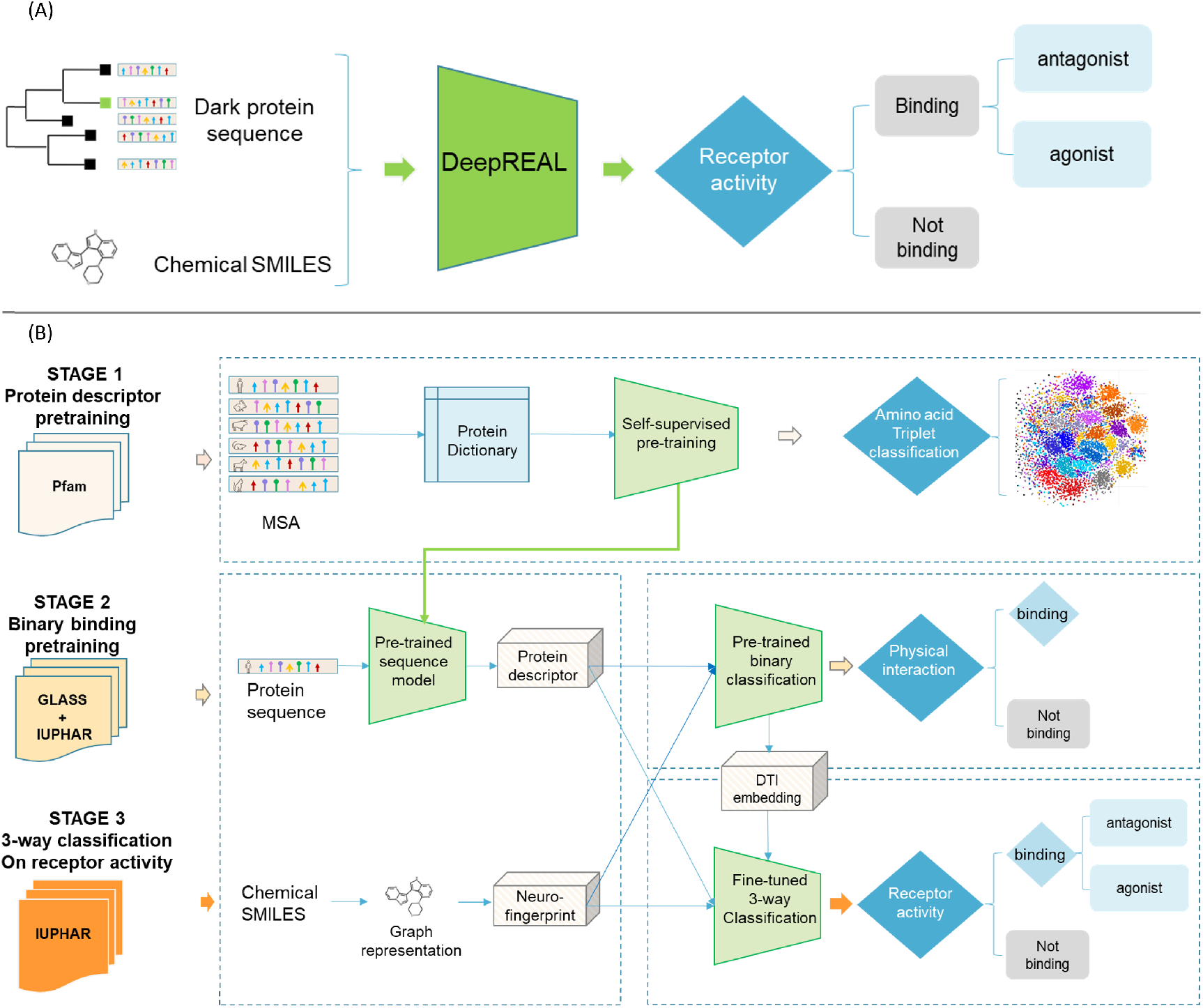
Illustration of DeepREAL. (A) Given a chemical and a protein sequence as inputs, DeepREAL will predict not only if the chemical is the ligand of protein but also the receptor activity upon the ligand binding. (B) DeepREAL is an end-to-end deep learning model trained using three stages. See text for details.

We collected receptor activity data from the International Union of Basic and Clinical Pharmacology/British Pharmacological Society (IUPHAR/BPS) Guide to Pharmacology and combined the additional Opioid data contributed in the paper [19]. There are 450 unique proteins involved in the complete receptor activity data set as shown in Table S1 in the Supporting Information.

To evaluate DeepREAL performance in light of real-world applications for dark proteins and novel chemicals, both data preprocessing and controlled experiment are designed to simulate various scenarios of data distribution shifts and to answer the following questions:

Q1: Is the pretraining helpful to improve the performance of receptor activity prediction using a small amount of data?
Q2: When DeepREAL is applied to unseen <u>dark proteins</u> that have low sequence similarity to those in the training data, what is the OOD generalization performance of DeepREAL?
Q3: When DeepREAL is used to predict unseen novel <u>chemicals</u> that are significantly different from those in the training data, what is the OOD generalization performance of DeepREAL?
Q4: When the test set label (agonist/antagonist/not-binding) distribution is close to reality and imbalanced compared to the training data, what is the generalization performance of DeepREAL?
Q5: How does DeepREAL perform compared to the state-of-the-art baseline models in both OOD and IID settings for predicting Opioid receptor activity?

### 2.2 Pretraining enables DeepREAL to generalize genome-scale receptor activity predictions using a relatively small data set

To answer Q1, the same model architecture is trained on the same IID and OOD settings using four procedures: 1) from total scratch without any pretraining, i.e., Stage 3 only, 2) going through Stage 1 whole Pfam pretraining but not the Stage 2 binary DTI classification pretraining, which is equivalent to the DISAE model[2], 3) going through only Stage 2 but not Stage 1, and 4) complete three stage pretraining/fine-tuning as DeepREAL. As shown in Figure 2 on the evaluation cross three classes (no-binding, agonist, antagonist), the model without any pretraining (i.e., only Stage 3) has the worst performance. Stage 1 or Stage 2 both boosts performance and the complete three-stage pipeline yields the best performance. From the by-class evaluation for antagonist or agonist as shown in Figure**??**, the precision and recall of DeepREAL is mostly higher than other variants in IID, protein OOD, and chemical OOD settings. Furthermore, the training curves of DeepREAL in Figure **??** converges faster than other variants in most cases. The advantage of pretraining is particularly apparent in chemical distribution shift OOD in the cross-class and the by-class evaluation as shown in Figure**??**, Figure 4 and Figure5. The chemical OOD is a more challenging OOD setting than other settings, where both chemical structure distribution and label ratio balance shift (more details in the following section). DISAE and the only-Stage 3 model have lower Cohen’s kappa, ROC-AUC, MCC than DeepREAL and the Stage2+Stage3 model. The later two model have relative close performance, suggesting that DTI pretraining plays a more important role the the sequence pretraining in the current training procedure. It may be because the whole-Pfam information learned at stage 1 is more difficult to transfer to Stage 3, as supported by the observation shown in Figure 4 and Figure5. It will be interesting to use other advanced training procedure such as prompting[6] or design different architecture (e.g., using skip connections[8][5] etc.).

**Figure 2:**
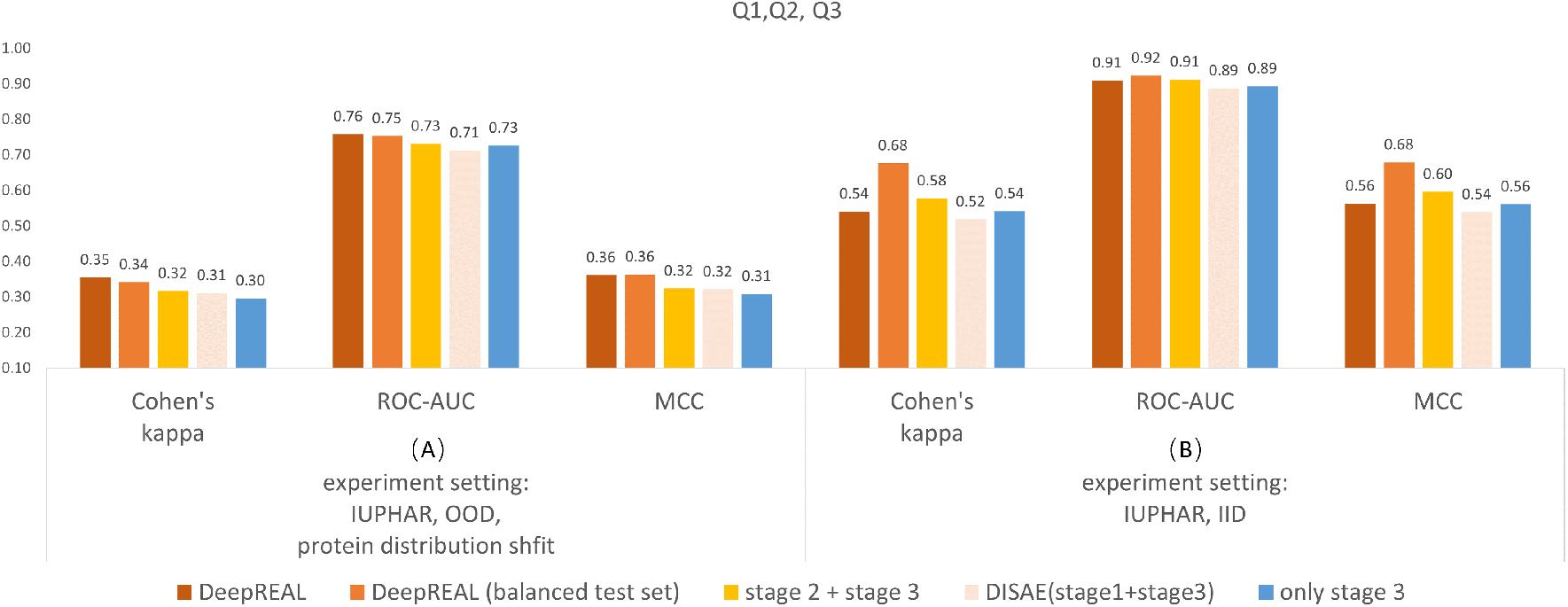
Performance comparison of DeepREAL with its variants in (A) protein OOD and (B) protein IID settings. The performance is evaluated by multiple gene families in the IUPHAR database.

**Figure 3:**
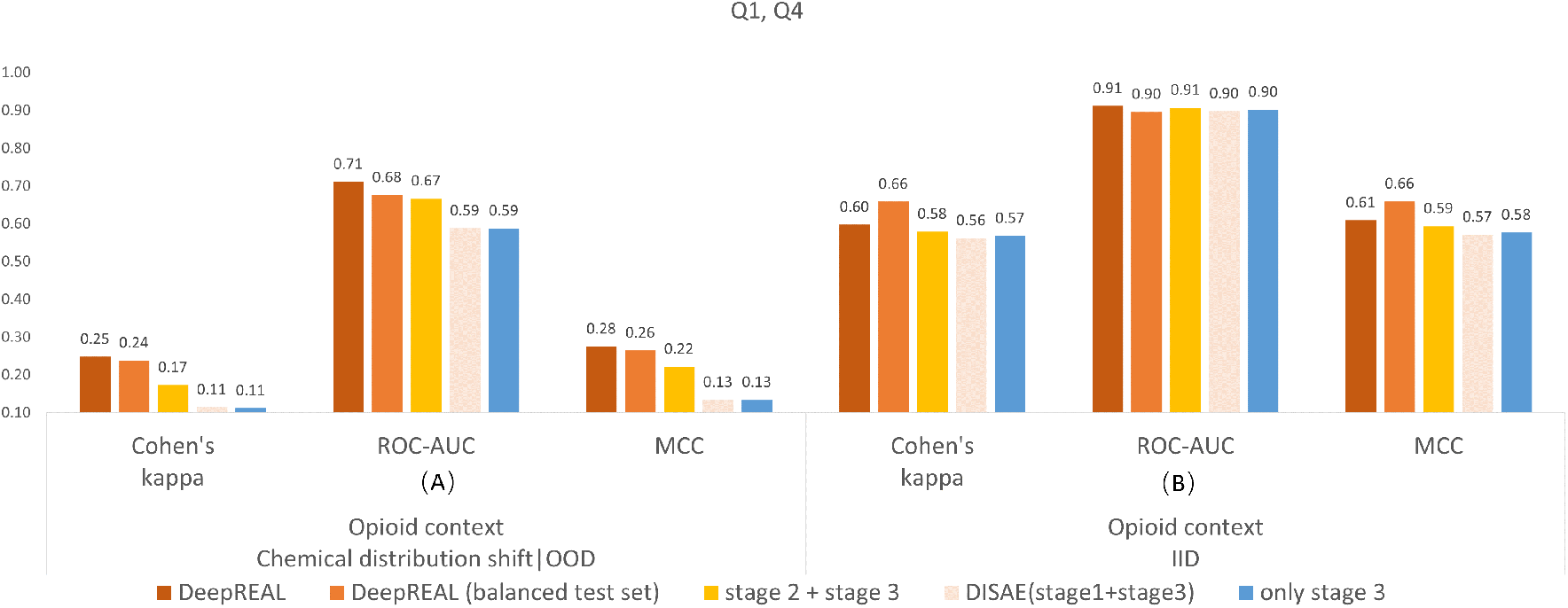
Performance comparison of DeepREAL with its variants in (A) chemical OOD and (B) chemical IID settings. The performance is evaluated only by opioid receptors.

**Figure 4:**
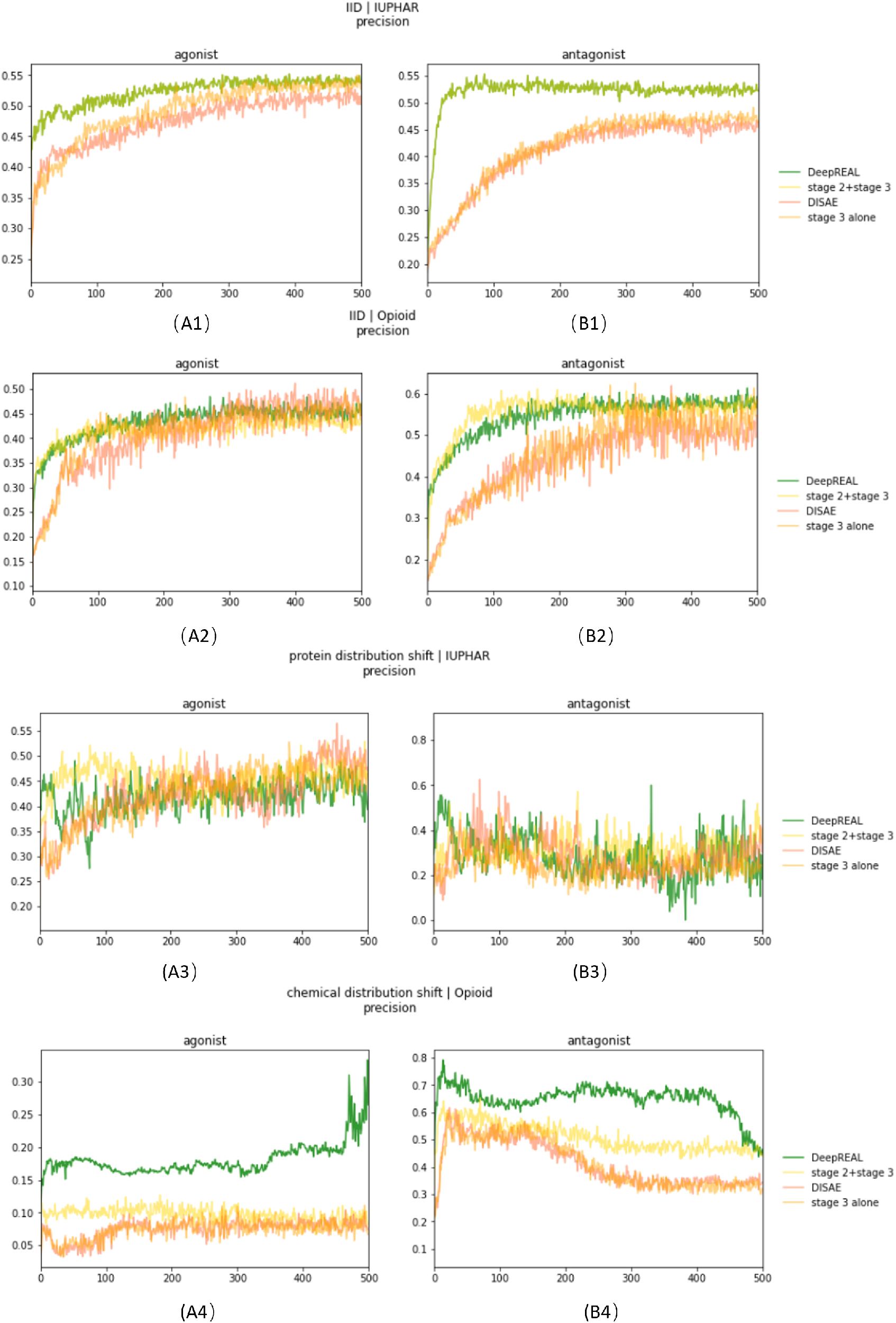
Training curves of DeepREAL and its variants when measured by the precision for predicting agonists or antagonists. The x-axis is number of training epochs. The y-axis is precision.

**Figure 5:**
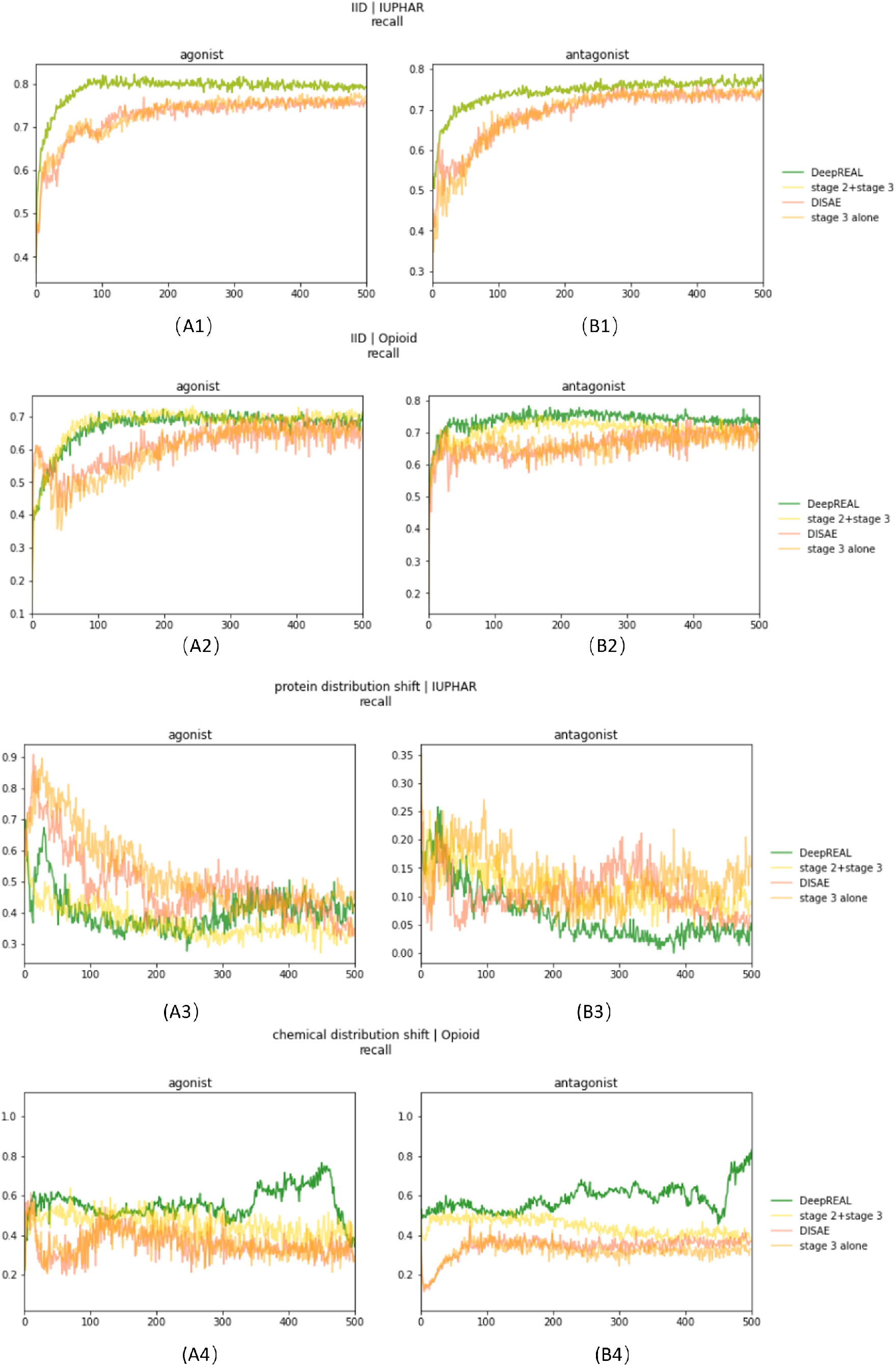
Training curves of DeepREAL and its variants when measured by the recall for predicting agonists or antagonists. The x-axis is number of training epochs. The y-axis is recall.

### 2.3 DeepREAL is robust in various shifted distribution scenarios

Q2, Q3, Q4 are three typical shifted distribution scenarios in real-world applications, i.e., the OOD generalization challenges. DeepREAL proves robust in each of the settings. It makes DeepREAL applicable to explore dark chemical genomics space.

Q2 focuses on the distribution shift coming from proteins. It is the dominant challenge when applying DeepREAL to a genome-scale given majority of proteins are dark without any receptor activity data. As shown in Figure**??**, by splitting the receptor activity data sets based on protein sequence similarities, 450 proteins and their associated interaction data are split to an OOD train/test setting. Under this setting, although the performance drops compared with the easier IID setting, the ROC-AUC score is still at 0.766 while existing state-of-the-art one-protein-one-model approach[19] is totally unable to make predictions in the protein OOD setting. The similarity distributions could be found in Supporting Figure S2.

In a similar fashion, by splitting the data based on the chemical similarity measured by Tanimoto coefficient, DeepREAL is evaluated in the setting of chemical distribution shift to answer Q3. Because the state-of-the-art model based on Random Forest[19] per protein is only applied to Opioid Receptors, only labeled data related to Opioid Receptors are included in the evaluation. The advantage of DeepREAL is apparent over other configurations including DISAE, as shown in Figure**??** Figure6A in the chemical OOD setting.

**Figure 6:**
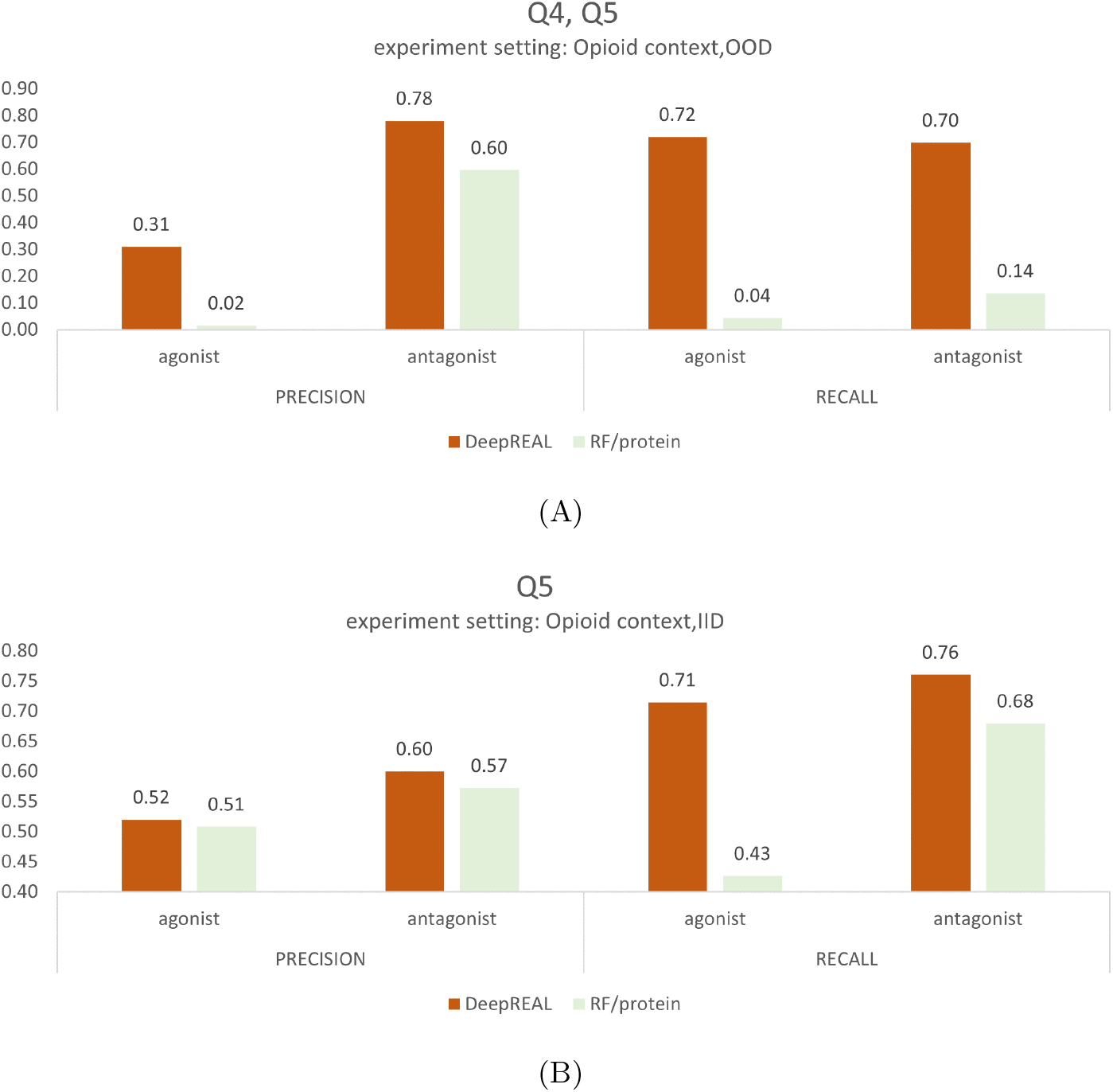
Performance comparison of DeepREAL with the state-of-the-art RF/protein model in (A) chemical OOD and, (B) chemical IID settings. The performance is evaluated by precision and recall for agonist or antagonist predictions

Although it is expected that a machine learning model performs the best when positive and negative data are balanced, the unseen binding/not-binding cases are imbalanced in reality, which has an estimated ratio of 1:5[12]. Hence, to answer Q4 about label distribution shift, for all experiments the number of not-binding samples in the test set is about five times as large as that of the agonist/antagonist data while training data are balanced for each class. For a comparison, a balanced test set is also evaluated. In general, DeepREAL evaluated by the balanced test set in the IID setting, which represents conventional cross-validations, outperforms that evaluated by the imbalanced data. However, in both protein and chemical OOD settings that simulates a real application, DeepREAL evaluated by the imbalanced data performs the best, as shown in Figure2 and Figure3. These observations suggest that the cross-validation in an IID setting is often over-optimistic and DeepREAL is more robust in a realistic application.

### 2.4 DeepREAL significantly outperforms state-of-the-art models

To compare DeepREAL with the leading machine learning model (RF/protein)[19] that can only predict opioid receptor activities, opioid receptor data set is split in two different ways for IID and OOD experiments as described in the previous section. In both IID and OOD settings, DeepREAL significantly outperforms the baseline in terms of precision and recall, as shown in Figure6A. Furthermore, the performance drop of DeepREAL from the IID setting to the OOD setting is less significant than that of the baseline.

To prove the statistical significance of DeepREAL performance against the baseline, the same training is repeated for five times under Opioid context with different random seeds. As shown in the Supporting Figure S3, the p-value of the hypothesis that the two models have the same average ROC-AUC is close to 0.0.

## 3 Conclusion

This paper proposed a deep learning framework DeepREAL that expands the traditional DTI task to predicting receptor activities of dark proteins and novel chemicals under various OOD settings. DeepREAL has several unique features. First, unlike the existing method that requires training one model for one protein and applying the trained model on the same protein, DeepREAL is needs only to train only one model to make predictions on any proteins with improved accuracy. Second, DeepREAL has improved generalization power in the face of all major types of deployment data distribution shifts, making it robust in real-world applications. Finally, by utilizing large unlabeled sequence data and rich binary bioassay data, DeepREAL models receptor activities on a multi-scale to alleviate data scarcity problem. Together, DeepREAL significantly outperforms existing algorithms for predicting receptor activities of ligand binding.

The performance of DeepREAL can be further improved along several directions. For example, unsupervised pretraining of chemical space could improve DeepREAL’s ability to detect novel chemicals[14]. Incorporating structure information into the protein sequence pretraining could help the downstream prediction tasks for ligand binding and receptor activity. We only predict two classes of receptor activity: agonist versus antigonist. In fact, the receptor activity is more complex than two mutually exclusive classes. There are other subtle activity classes such as partial agonist. A multi-class model could be a more suitable choice and subject to future studies.

## 4 Methods

### 4.1 Data

There are four data sets involved: Pfam used to pretrain protein descriptor [17]; GLASS data [3] with a large GPCR protein-ligand binding binary data set; agonist/antagonist data are downloaded from the International Union of Basic and Clinical Pharmacology/British Pharmacological Society (IUPHAR/BPS) Guide to Pharmacology; Opioid receptor activity data from [19]. Protein descriptor pretraining exactly follows DISAE[2] hence is not explained in details here. In brief, DISAE builds up a distilled triplet sequence dictionary for the whole Pfam proteins based on multiple sequence alignments (MSA). Every input protein will be mapped to its distilled triplets representation according to the protein dictionary as illustrated in 1. Chemical-protein pairs with the receptor activity annotation is treated as positive in the binary DTI setting and combined with GLASS for the binary classification pretraining. IUPHAR/BPS combined with [19] Opioid data makes the final data set used in Stage3 three-way classification. Detailed data statistics could be found Supporting information Table S1.

In terms of data splitting, IID setting splits the data randomly as conventional cross-validations, except for the Opioid context experiments where all three Opioid proteins are ensured to appear in both training and testing data sets. The OOD data split is carried out with a spectral clustering algorithm[15] based on pair-wise similarity scores. The similarity distributions could be found in Supporting Figure S2. In our experiments, the Stage 2 binary training is always carried out with the same data.

### 4.2 Baseline

To compare with baseline[19] which showed Random Forest model with PubChem fingerprints^1^ [1] had the best performance, the same fingerprint feature was generated for all Opioid paired chemicals to train Random Forest models. The baseline was chosen for it being, as far as we know, the first and only work in the direction of receptor activity prediction. Keeping other hyper-parameters the same as in [19], the Random Forest depth is tuned to find the best performance model for each Opioid protein with an example curve in Supporting Figure S4. For each experiment, one Random Forest is trained for each Opioid protein. An average Random Forest test performance is calculated weighted by the sample size of each Opioid protein. When evaluating the model performance variance, different random seeds were used.

### 4.3 DeepREAL framework

#### 4.3.1 Architecture

DeepREAL has a novel three-stage framework. There are four major modules in DeepREAL model: protein sequence embedding, chemical structure descriptor, binary interaction learner and multi-class classifier as shown in Supporting Figure S1. Under this framework, the state-of-the-art binary DTI model DISAE[2] was employed as model architecture backbone, which includes ALBERT[11] based protein descriptor, and attentive pooling[20] based binary interaction learner. Built on DISAE, the chemical descriptor architecture in DeepREAL is changed to the state-of-the-art graph neural network ContextPred[9].

The new component multi-class classifier includes two sub-modules: a three-way interaction learner uses the same architecture as the binary interaction learner; after concatenating all related embeddings, the concatenated tensor goes through a ResNET[8] layer and MLP[7] transformation to generate the final logit vector used in cross entropy loss calculation.

### 4.3.2 Framework information flow

The knowledge transfer across stages was realized by sharing weights on the first three modules in DeepREAL architecture, i.e., protein descriptor, chemical descriptor and binary interaction learner. Protein descriptor first went through Stage 1 whole Pfam pretraining in self-supervised fashion, transferring whole Pfam protein syntax to the Stage 2 binary pretraining, and then, together with newly initialized chemical descriptor and binary interaction learner, learns to predict whether or not a protein and chemical would bind on GLASS data in a supervised learning manner. Weights of these three modules would all be transferred to Stage 3.

In Stage 3, the three modules were first duplicated: one copy had frozen weights whereas the other copy updated its weights for *n* epochs with multi-class-learner on DeepREAL receptor activity information in a supervised learning manner, where *n* is a hyperparameter as shown in Supporting Information Figure S1. In our experiments, we fould that n being small, such as 50, would help to improve model generalization performance when the training data size was smaller. This phenomenon is due to the fact that this complete model with so many trainable parameters is capable of memorizing a small training set, resulting in over-fitting and poor generalization. More frozen weights would limit the over-fitting and put more pressure on the multi-class-learner to learn robust representation.

The protein embedding vector, chemical embedding vector, and binary interaction embedding vector output by one copy of the binary pretrained modules went into three-way interaction learner to learn a three-way interaction embedding. Output embedding from all the component, the seven embedding vectors were concatenated and fed into ResNET[8] and MLP[7] to make the final three-way classification, after which the information flow within DeepREAL was completed.

Three stage model was designed to train sequentially and separately. Only optimized weights were transferred. The Stage 1 optimization procedure has been described in [2]. Stage 2 and Stage 3 optimization were both driven by cross-entropy loss in a stochastic manner using Adam[10].

#### 4.3.3 Pretraining implementation and module frozen strategy

A key element of success in multi-stage pretraining is to transfer knowledge. A major challenge in the three-stage pipeline is to prevent the previously learned knowledge from being lost during the weight update in the subsequent stage. DISAE has reported the benefits of a frozen mechanism. This strategy was adopted in DeepREAL Stages 2 and 3 as well. In Stage 2, following the experience of DISAE, part of the transformer[21] layers were frozen. In Stage 3, the binary pretrained modules were duplicated to have one copy always frozen and the other copy fine-tuned for only *n* epochs. Without tuning, *n* was empirically set to 50 in the Opioid receptor focused experiments while on the complete DeepREAL receptor data set involving 450 proteins, *n* was set as infinity until the model converges.

## Acknowledgement

This project has been funded with federal funds from the National Institute of General Medical Sciences of National Institute of Health (R01GM122845) and the National Institute on Aging of the National Institute of Health (R01AD057555).

## Author Contributions

LX conceived and planned the experiments. TC developed and implemented the algorithm. TC, KAA, and YL carried out the experiments. LX, TC contributed to the interpretation of the results. TC, KAA, YL and LX wrote the manuscript. All authors provided critical feedback and helped shape the research, analysis and manuscript.

## Footnotes

1 ftp://ftp.ncbi.nlm.nih.-gov/pubchem/specifications/pubchem_fingerprints.txt

## References

[1] Evan E Bolton, Yanli Wang, Paul A Thiessen, and Stephen H Bryant. Pubchem: integrated platform of small molecules and biological activities. In Annual reports in computational chemistry, volume 4, pages 217–241. Elsevier, 2008.

[2] Tian Cai, Hansaim Lim, Kyra Alyssa Abbu, Yue Qiu, Ruth Nussinov, and Lei Xie. Genome-wide prediction of small molecule binding to remote orphan proteins using distilled sequence alignment embedding. bioRxiv, 2020.

[3] Wallace K. B. Chan, Hongjiu Zhang, Jianyi Yang, Jeffrey R. Brender, Junguk Hur, Arzucan Özgür, and Yang Zhang. GLASS: a comprehensive database for experimentally validated GPCR-ligand associations. Bioinformatics, 31(18):3035–3042, 05 2015.

[4] Joseph A DiMasi, Henry G Grabowski, and Ronald W Hansen. Innovation in the pharmaceutical industry: new estimates of r&d costs. Journal of health economics, 47:20–33, 2016.

[5] Alexey Dosovitskiy, Lucas Beyer, Alexander Kolesnikov, Dirk Weissenborn, Xiaohua Zhai, Thomas Unterthiner, Mostafa Dehghani, Matthias Minderer, Georg Heigold, Sylvain Gelly, et al. An image is worth 16×16 words: Transformers for image recognition at scale. arXiv preprint arXiv:2010.11929, 2020.

[6] Tianyu Gao, Adam Fisch, and Danqi Chen. Making pre-trained language models better few-shot learners. arXiv preprint arXiv:2012.15723, 2020.

[7] Trevor Hastie, Robert Tibshirani, and Martin Wainwright. Statistical learning with sparsity: the lasso and generalizations. Chapman and Hall/CRC, 2019.

[8] Kaiming He, Xiangyu Zhang, Shaoqing Ren, and Jian Sun. Deep residual learning for image recognition. CoRR, abs/1512.03385, 2015.

[9] Weihua Hu, Bowen Liu, Joseph Gomes, Marinka Zitnik, Percy Liang, Vijay Pande, and Jure Leskovec. Strategies for pre-training graph neural networks. arXiv preprint arXiv:1905.12265, 2019.

[10] Diederik P Kingma and Jimmy Ba. Adam: A method for stochastic optimization. arXiv preprint arXiv:1412.6980, 2014.

[11] Zhenzhong Lan, Mingda Chen, Sebastian Goodman, Kevin Gimpel, Piyush Sharma, and Radu Soricut. Albert: A lite bert for self-supervised learning of language representations. arXiv preprint arXiv:1909.11942, 2019.

[12] Hansaim Lim, Aleksandar Poleksic, Yuan Yao, Hanghang Tong, Di He, Luke Zhuang, Patrick Meng, and Lei Xie. Large-scale off-target identification using fast and accurate dual regularized one-class collaborative filtering and its application to drug repurposing. PLoS computational biology, 12(10):e1005135, 2016.

[13] Ann Lin, Christopher J Giuliano, Ann Palladino, Kristen M John, Connor Abramowicz, Monet Lou Yuan, Erin L Sausville, Devon A Lukow, Luwei Liu, Alexander R Chait, et al. Off-target toxicity is a common mechanism of action of cancer drugs undergoing clinical trials. Science translational medicine, 11(509), 2019.

[14] Yang Liu, You Wu, Xiaoke Shen, and Lei Xie. Covid-19 multi-targeted drug repurposing using few-shot learning. Frontiers in Bioinformatics, 1:18, 2021.

[15] Ulrike Von Luxburg. A tutorial on spectral clustering, 2007.

[16] James J III Lynch, Terry R Van Vleet, Scott W Mittelstadt, and Eric AG Blomme. Potential functional and pathological side effects related to off-target pharmacological activity. Journal of pharmacological and toxicological methods, 87:108–126, 2017.

[17] Jaina Mistry, Sara Chuguransky, Lowri Williams, Matloob Qureshi, Gustavo A Salazar, Erik LL Sonnhammer, Silvio CE Tosatto, Lisanna Paladin, Shriya Raj, Lorna J Richardson, et al. Pfam: The protein families database in 2021. Nucleic Acids Research, 49(D1):D412–D419, 2021.

[18] Tudor I Oprea. Exploring the dark genome: implications for precision medicine. Mammalian Genome, 30(7):192–200, 2019.

[19] Srilatha Sakamuru, Jinghua Zhao, Menghang Xia, Huixiao Hong, Anton Simeonov, Iosif Vaisman, and Ruili Huang. Predictive models to identify small molecule activators and inhibitors of opioid receptors. Journal of Chemical Information and Modeling, 2021.

[20] Cicero dos Santos, Ming Tan, Bing Xiang, and Bowen Zhou. Attentive pooling networks. arXiv preprint arXiv:1602.03609, 2016.

[21] Ashish Vaswani, Noam Shazeer, Niki Parmar, Jakob Uszkoreit, Llion Jones, Aidan N Gomez, Łukasz Kaiser, and Illia Polosukhin. Attention is all you need. In Advances in neural information processing systems, pages 5998–6008, 2017.

[22] Chi Heem Wong, Kien Wei Siah, and Andrew W Lo. Estimation of clinical trial success rates and related parameters. Biostatistics, 20(2):273–286, 2019.

[23] Lei Xie, Li Xie, Sarah L Kinnings, and Philip E Bourne. Novel computational approaches to polypharmacology as a means to define responses to individual drugs. Annual review of pharmacology and toxicology, 52:361–379, 2012.

