## Supplementary Information for "DeepREAL: A Deep Learning Powered Multi-scale Modeling Framework Towards Predicting Out-of-distribution Receptor Activity of Ligand Binding"

### Supporting Information

September 13, 2021

| | unique protein | unique chemical | agonist-protein pair | antagonist-protein pair | not-binding chemical-protein pair |
|---|---|---|---|---|---|
| IUPHAR | 450 | 13,126 | 14,412 | 14,488 | 144,500 |
| Opioid receptors related | 3 | 2,483 | 2,920 | 2,996 | 29,580 |

Table S 1: Data statistics of the 3-way classification data sets.

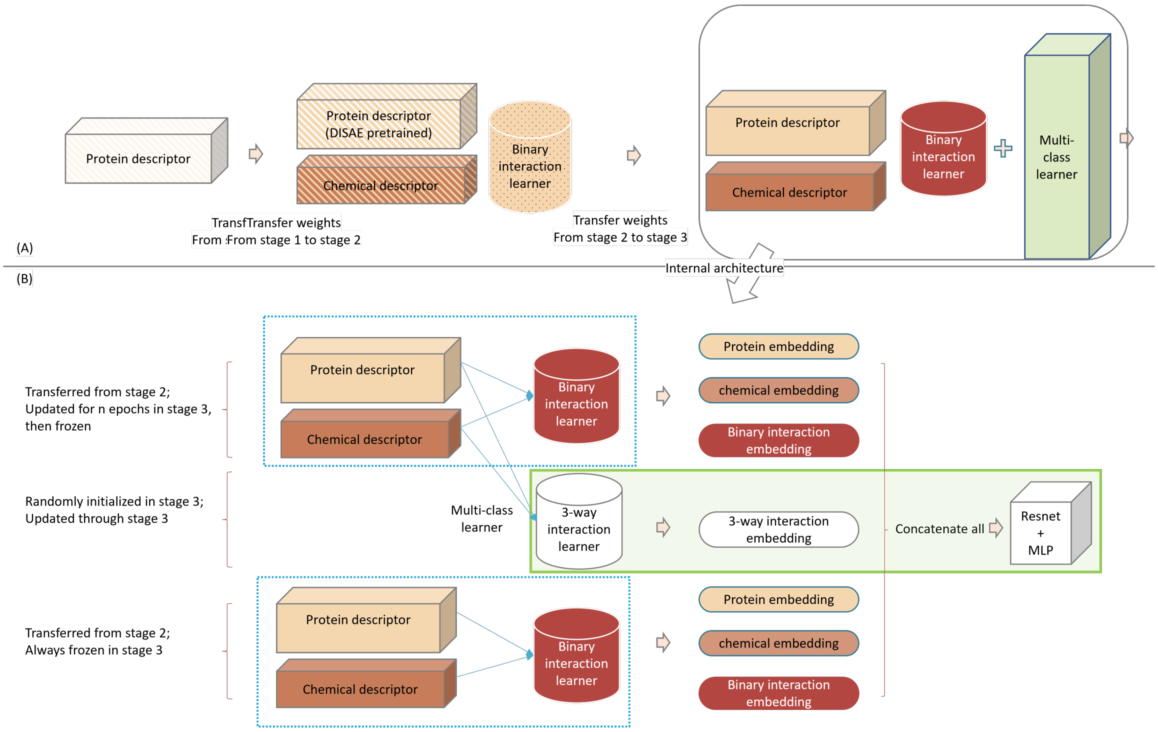


Figure S 1: The architecture details and information flow of DeepREAL. (A) shows the transfer learning across three stages.(B) shows the internal architecture at stage 3.

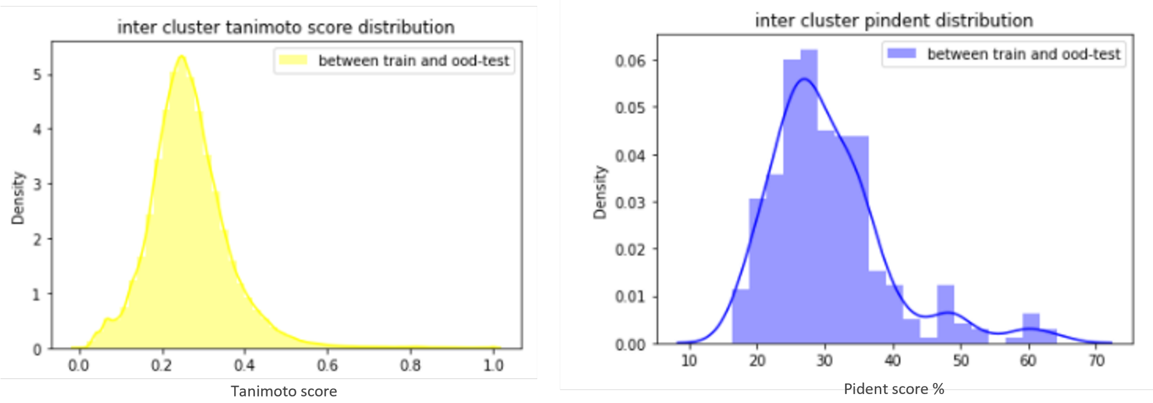


Figure S 2: OOD data split controls similarity between training data and OOD test data to simulate real-life deployment data distribution shift. Tanimoto score is used to measure chemical similarity and the left panel shows the distribution of pair-wise Tanimoto scores between chemicals in the train set and chemicals in the test, showing that chemicals in OOD-test set is significantly different from chemicals in the training set. In the same fashion, the pident score is used to measure protein similarity. The right panel shows the distribution of pair-wise pident scores between proteins in the training set and proteins in the test set.

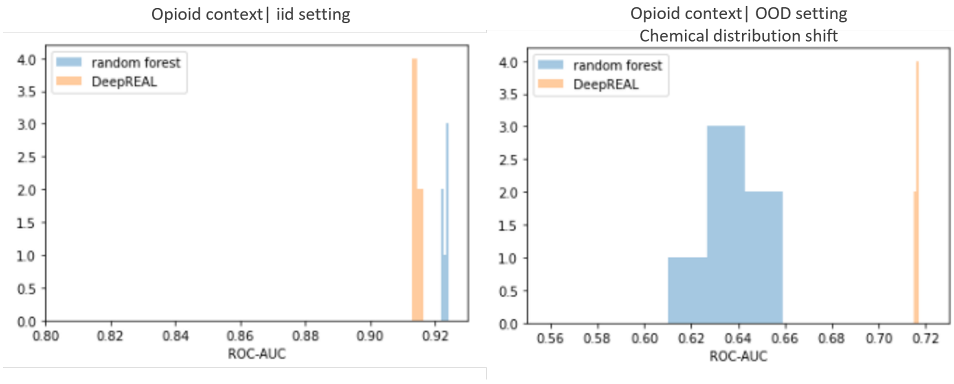


Figure S 3: ROC-AUC score distribution under Opioid context when repeating the experiments five times with different random seeds. On the left panel, the experiment data is IID random split. In this scenario, although Random Forest has statistical significant better performance, the margin between the mean ROC-AUC scores is only 0.006. On the right panel, the experiment data is OOD split to simulate deployment data chemical distribution shift. In this scenario, DeepREAL not only shows statistically better ROC-AUC with smaller standard deviation but also has 0.08 margin in terms of mean ROC-AUC.

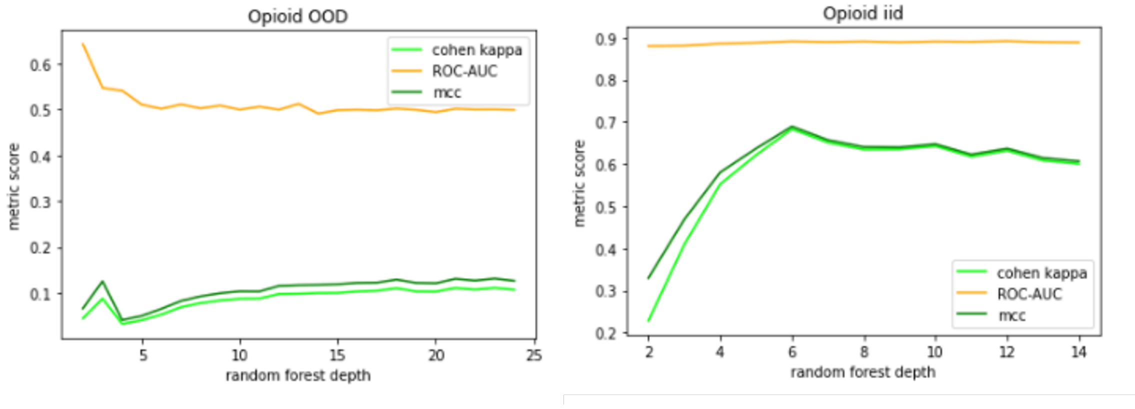


Figure S 4: Random Forest models are tuned to find the best depth while keeping all other hyperparameters the same as in the baseline work. Here's an example of tuning for one Opioid 'P35372' receptor's Random Forest models under both IID and OOD settings. The same tuning is done for all other two proteins as well, and the best model for each protein is used to make the test set prediction evaluation.
